## Supplementary figures and images for "Re-exposure to a sensorimotor perturbation produces opposite effects on explicit and implicit learning processes"

### Supplemental Figure 1

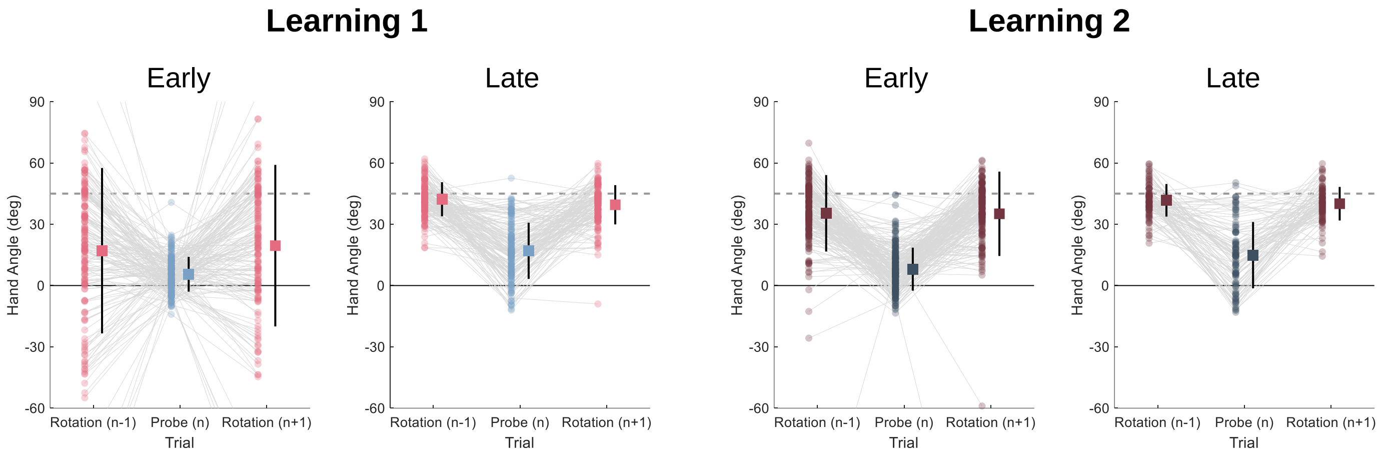

### Supplemental Figure 2

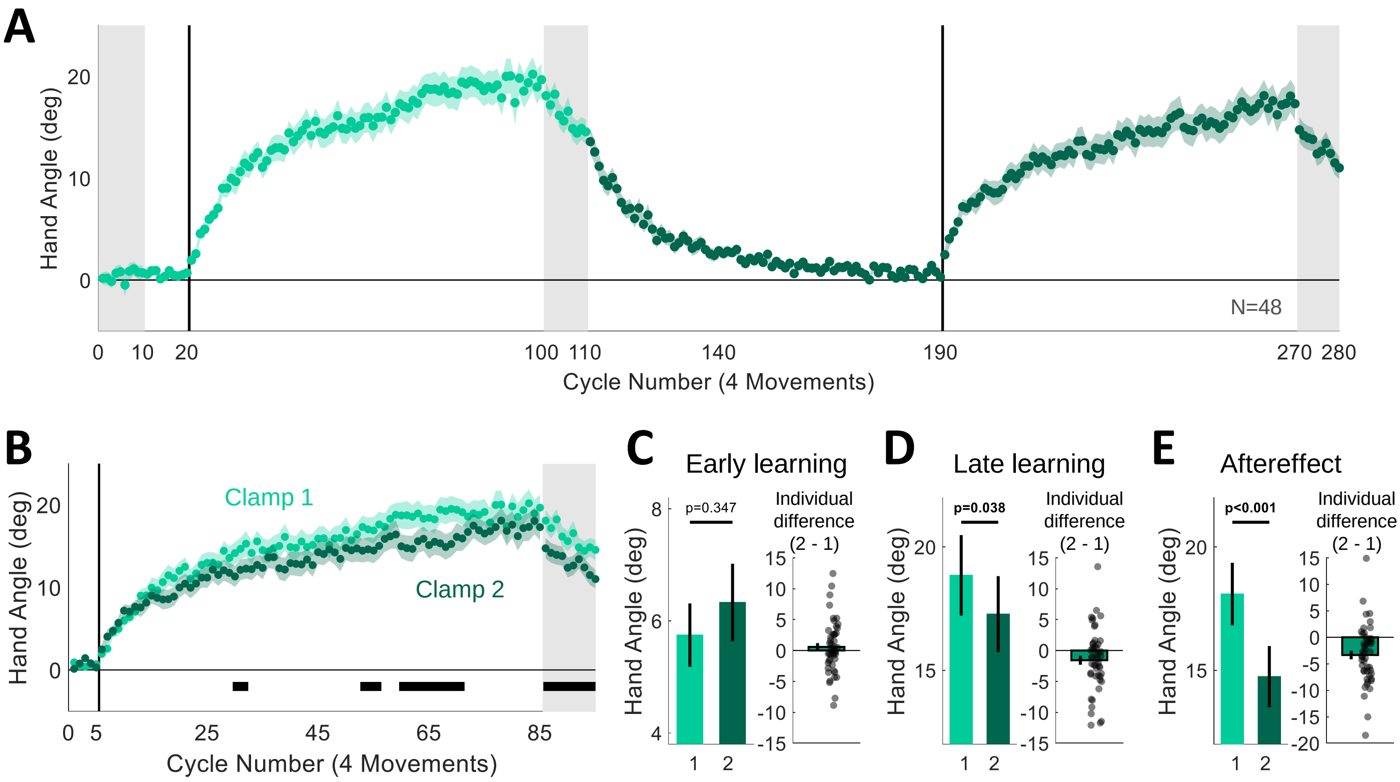
